# Genome size and the extinction of small populations

**DOI:** 10.1101/173690

**Authors:** Thomas LaBar, Christoph Adami

## Abstract

Although extinction is ubiquitous throughout the history of life, insight into the factors that drive extinction events are often difficult to decipher. Most studies of extinction focus on inferring causal factors from past extinction events, but these studies are constrained by our inability to observe extinction events as they occur. Here, we use digital evolution to avoid these constraints and study “extinction in action”. We focus on the role of genome size in driving population extinction, as previous work both in comparative genomics and digital evolution has shown a correlation between genome size and extinction. We find that extinctions in small populations are caused by large genome size. This relationship between genome size and extinction is due to two genetic mechanisms that increase a population’s lethal mutational burden: large genome size leads to both an increased lethal mutation rate and an increased likelihood of stochastic reproduction errors and non-viability. We further show that this increased lethal mutational burden is directly due to genome expansions, as opposed to subsequent adaptation after genome expansion. These findings suggest that large genome size can enhance the extinction likelihood of small populations and may inform which natural populations are at an increased risk of extinction.

## Introduction

The ubiquity of extinction events throughout the history of life [1], and the increasing realization that the biosphere may be experiencing a sixth mass extinction [2] drives interest in determining the factors that cause certain species, but not others, to go extinct [3]. It is accepted that a combination of genetic [4, 5], demographic [6, 7], environmental [8, 9], and ecological [10, 11, 12] factors contribute to species extinctions. Beyond those deterministic factors, chance events also likely influence certain extinction events [13, 14]. Here, we focus on the genetic factors influencing extinction, specifically the role of small population size and genetic drift [15].

In small populations, weakened purifying selection leads to increased fixation of small-effect deleterious mutations [16]. As multiple deleterious mutations fix, the absolute fitness of the population may decrease, resulting in a decrease in population size. This decreased population size further weakens selection, leading to the fixation of additional deleterious mutations and a further decrease in population size. This process continues until the population goes extinct. This positive feedback loop between decreased population size and deleterious mutation fixation is known as a mutational meltdown [17]. Mathematical models of mutational meltdowns suggest that even intermediate-sized asexual populations can quickly go extinct [18, 19]. Likewise, small sexual populations are also vulnerable to fast meltdowns [20]. The detection of mutational meltdowns in natural populations is difficult; other non-genetic factors (i.e., environmental or ecological factors) can obscure the role of mutation accumulation in extinction.

While the concept of a mutational meltdown provides a population-genetic mechanism for extinction, it is still uncertain what factors beyond population size influence the likelihood of a meltdown. For example, if deleterious mutation accumulation drives mutational meltdowns, then species with a greater genomic mutation rate should be at a greater risk of extinction [21, 22]. Therefore, genetic mechanisms that increase the genomic mutation rate may also increase the likelihood of species extinction. One such genetic mechanism that could increase the mutation rate are genome expansions (i.e., mutations that increase genome size) because species with larger genomes (but similar point mutation rates) have greater genomic mutation rates. If these genome expansions lead to an increase in functional genome content, then the deleterious mutational load of a population should also increase. Indeed, there is some evidence that genome size positively correlates with extinction risk in certain clades of multicellular organisms [23, 24].

It is difficult to experimentally test the role of genome size in extinction in both natural and laboratory model systems. Here, we use digital experimental evolution to test whether genome expansions can drive population extinction. Digital experimental evolution is the computational counterpart to microbial experimental evolution [25, 26, 27]. Instead of a population of microbes evolving in a flask (or other physical microcosm), digital evolution experiments instantiate a population of self-replicating computer programs that reproduce and mutate in a digital world [28]. Most digital evolution systems do not try to emulate any specific biological system. Instead, these systems implement populations with heritability, variation, and differential fitness (i.e., the three requirements for Darwinian evolution) but composed of organisms and genomes significantly simpler than those in biological populations [29].

In a previous study with the digital evolution system Avida [30] on the role of population size in the evolution of complexity, we found that the smallest populations evolved the largest genomes and the most novel traits, but also had the greatest extinction rates [31]. Here, we use Avida to test explicitly the role of genome size in the extinction of small populations. Avida differs from previous models of extinction in small populations in the mode of selection. Unlike mutational meltdown models [15], where selection is hard and the accumulation of deleterious mutations directly leads to population extinction, selection is primarily soft in Avida and deleterious mutations alter relative fitness (i.e., competitive differences between genotypes), not absolute fitness (i.e., differences in the number of viable offspring between genotypes). Extinction occurs in Avida through the accumulation of “lethal” mutations, or more precisely, “non-viable” mutations that prevent their bearer from reproducing. These non-viable avidians occupy a portion of the limited space allocated to an avidian population, thus reducing the effective population size and potentially causing extinction over time.

We find that extinction in small populations is to a large extent driven by genome expansions, and that an increased genome size not only leads to an increase in the genomic mutation rate, but specifically to an increase in the lethal mutation rate. Heightened lethal mutation rates eventually lead to population extinction. Additionally, we show that genotypes with large genomes have an elevated probability of stochastic replication errors during reproduction (i.e., stochastic viability), further elevating the likelihood of non-viability of offspring and extinction. These results suggest that large genome size does elevate the risk of population extinction due to an increased lethal mutational burden.

## Methods

### Avida

For the following experiments, we used the digital experimental evolution platform Avida, version 2.14 [30]. In Avida, simple computer programs (“avidians”) compete for the resources required to undergo self-replication and reproduction. Each avidian consists of a genome of computer instructions drawn from a set of twenty-six available instructions in the Avida genetic code. A viable asexual avidian genome must contain the instructions to allocate a new (offspring) avidian genome, copy the instructions from the parent genome to the offspring genome, and divide off the offspring genome into a new avidian. During this copying process, mutations may occur that introduce variation into the population. These novel mutations can then be passed onto future generations, resulting in heritable variation. This genetic variation causes phenotypic variation: avidians with different genomes may self-replicate at different speeds. As faster self-replicators will outcompete slower self-replicators, there is differential fitness between avidians. Therefore, given there is heritable variation and differential fitness, an Avida population undergoes Darwinian evolution [32, 29]. Avida has previously been used to test hypotheses concerning the evolution of genome size [33, 31], the role of population size in evolution [34, 35, 31, 36], and the consequences of population extinction [37, 38, 39, 40].

The Avida world consists of a grid of *N* cells; each cell can be occupied by at most one avidian. Thus, *N* is the maximum population size for the Avida environment. While Avidian populations are usually at carrying capacity, the presence of lethal mutations can reduce their effective population size below this maximum size. In traditional Avida experiments, the geometry of the environment can alter evolutionary dynamics, as offspring are placed into the environment in one of nine cells neighboring (and including) their parent’s cell [41]. Here, offspring can be placed into any cell in the environment, simulating a well-mixed environment (i.e., no spatial structure). If a cell is occupied by another avidian, the new offspring will overwrite the occupant. The random placement of offspring avidians adds genetic drift to Avida populations, as avidians are overwritten without regard to fitness.

Time in Avida is structured in discrete units called “updates”. During each update, 30N genome instructions are executed across the population. The ability for an avidian to execute one instruction in its genome is called a SIP, or Single Instruction Processing unit. In a population consisting of *N* individuals with the same genotype, each avidian will receive approximately 30 SIPs, and thus execute 30 instructions each update. However, in a population with multiple genotypes, some genotypes may be allocated more SIPs than others, depending on a value called “merit”; genotypes with greater merit will receive proportionally more SIPs than genotypes with lesser merit.

An avidian lineage can evolve increased merit through two means. First, an increase in genome size will increase merit by a proportional amount. Merit is set to be proportional to genome size in order to offset the decrease in replication speed, and thus the decrease in fitness, caused by increasing genome size. The second way to increase merit is through the evolution of certain phenotypic traits. Avidians can evolve the ability to perform Boolean logic calculations. If an avidian can input random numbers from the environment, perform a calculation using these numbers and output a correct result, its offspring’s merit will be increased by a preset amount. The performance of calculations of greater complexity will result in a greater merit improvement. In the experiments here that select for trait evolution, we used the so-called “Logic-9” environment [41]. In this environment, the performance of NOT or NAND multiplies merit by 2, the performance of ORNOT or AND multiplies merit by 4, the performance of OR or AND NOT multiplies merit by 8, the performance of NOR or XOR multiplies merit by 16, and the performance of EQUALS multiplies merit by 32. If a genotype can perform multiple calculations, the merit multiplier is multiplicative (i.e, a genotype that can perform NOT and NAND for example has its merit multiplied by 4).

Fitness for an avidian genotype is estimated as the genotype’s merit divided by its gestation time (the number of instruction executions needed for reproduction). Thus, fitness is the ratio of the number of instructions a genotype can execute in a given time to the number of instructions it needs to execute to reproduce. Therefore, there are two avenues for a population of avidians to increase fitness: increase their merit or decrease the number of instruction executions needed for self-replication.

There are a variety of different possible implementations of mutations in Avida. Here, we used settings that differed from the default in order to improve our ability to analyze the causes of population extinction (see Table 1 for a listing of changes to the default settings). Point mutations occur upon division between parent and offspring, after replication. There is an equal probability that each instruction in the genome will receive a point mutation upon division; thus, genome size determines the total genomic mutation rate. To model indels, we used so-called “slip” mutations. This mutational type will randomly select two loci in the genome and then, with equal probability, either duplicate or delete the section of the genome between those two loci. Finally, to to ease our analysis, we required every offspring genotype to be equivalent to its parent’s genotype before the above mutations were applied at division.

In Avida, it is possible to perform experiments where the appearance of mutations with certain effects is prevented [42]. For example, it is possible to revert a mutation of a particular predetermined size after it has appeared. For this to occur, the Avida program analyzes the fitness of every novel genotypes that enters the population and, if the fitness is of the pre-set effect, the mutation is reverted. This system allows experimenters the ability to determine the relevance of certain mutational effects to evolution. However, mutations of certain effects can still enter the population if their fitness effect is stochastic. An avidian has stochastic fitness if its replication speed depends on characteristics of the random numbers it inputs in order to do its Boolean logic calculations. Some stored numbers may alter the order in which certain instructions are executed or copied into an offspring’s genome, thus altering fitness.

**Table 1:** Notable Avida parameters changed from default value.

| Parameter | Default Value | Changed Value | Treatment |
| --- | --- | --- | --- |
| WORLD_X | 60 | $N$ | All |
| WORLD_Y | 60 | 1 | All |
| BIRTH_METHOD | 0 | 4 | All |
| COPY_MUT_PROB | 0.0075 | 0.0 | All |
| DIV_MUT_PROB | 0.0 | $\mu$ | All |
| DIVIDE_INS_PROB | 0.05 | 0.0 | All |
| DIVIDE_DEL_PROB | 0.05 | 0.0 | All |
| DIVIDE_SLIP_PROB | 0.0 | 0.01 | All Variable Genome Size |
| REQUIRE_EXACT_COPY | 0 | 1 | All |
| REVERT_DEAD | 0.0 | 1.0 | Lethal-reversion |
| REVERT_DETRIMENTAL | 0.0 | 1.0 | Deleterious-reversion |

### Experimental Design

To study the role of genome size in the extinction of small populations, we first evolved populations across a range of per-site mutation rates (*μ* = 0.01 and *μ* = 0.1) and population sizes (*N* = {5,6, 7, 8,10,15, 20} for *μ* = 0.01 and *N* = {10,12,15,16,17, 20, 25} for *μ* = 0.01). For each combination of population size and mutation we evolved 100 populations for 10^5^ generations. Each population was initialized at carrying capacity with *N* copies of the default Avida ancestor with all excess instructions removed; this resulted in an ancestor with a genome of 15 instructions (only those needed for replication). Ancestral genotypes with persite mutation rates of *μ* = 0.01 and *μ* = 0.1 thus have genomic mutation rates of *U* = 0.15 and *U* =1.5 mutations/genome/generation, respectively. Genome size mutations *(indels) occur at a rate of 0.01 mutations/genome/generation for all treatments. Additionally, for each mutation rate and population size combination, an additional 100 populations were evolved in an environment where genome size was fixed. To directly test for the role of lethal and deleterious mutations in driving extinction, we evolved 100 populations at the low mutation rate population sizes under conditions where either lethal mutations or deleterious, but non-lethal, mutations were reverted (the “lethal-reversion” and “deleterious-reversion treatments”, respectively).

### Data Analysis

For all evolution experiments, we saved data on the most abundant (dominant) genotype every ten generations. The final saved dominant genotype was used in all analyses here. All data represents either genotypes at most ten generations before extinction (in the case of extinct populations) or genotypes from the end of the experiment (in the case of surviving populations). In order to calculate the lethal mutation rate and other relevant statistics for a genotype, we generated every single point mutation for that genotype and measured these mutants’ fitness using Avida’s Analyze mode. The lethal mutation rate was estimated as *U*_Lethal_ = *μ* × *L* × *p*_Lethal_, where *μ* is the per-site mutation rate, *L* is the genome size, and *P*_Lethal_ is the probability that a random mutation will be lethal.

### Analysis of the relationship between genome expansions and changes in the lethal mutation rate

To test whether genome expansions themselves were responsible for the increase in the lethal mutation rate or whether the lethal mutation rate increased after adaptation occurred in response to a genome expansion, we first reconstructed the line-of-descents (LODs) for each of the one hundred genotypes evolved in a population of 20 individuals with a per-site mutation rate of 0.01 mutations/site/generation. An LOD contains every intermediate genotype from the ancestral genotype to an evolved genotype and allows us to trace how genome size evolved over the course of the experiment [41]. We reduced these LOD to only contain the ancestral genotype, the genotypes that changed in genome size, the genotype immediately preceding a change in genome size, and the final genotype. We measured the genome size and the lethal mutation rate for each of these remaining genotypes. Then, we measured the relationship between the change in genome size and the change in the lethal mutation rate for genome expansions, genome reductions, and the segments of evolutionary time where genome size was constant.

### Analysis of stochastic viability

In order to test the possibility that some of our populations had evolved stochastic viability, we analyzed each genotype from the *N* = 5 lethal-reversion populations and each genotype from the *N* = 8, *μ* = 0.01, original populations. These population sizes were chosen because they had the greatest equality between number of extinct populations and number of surviving population for their respective treatments. We performed 100 viability trials, where a genotype was declared non-viable if it had a fitness equal to 0. A genotype was declared stochastic-viable if the number of trials where it was nonviable was greater than 0 and less than 100. Otherwise, it was defined as deterministic-viable or always viable.

All data analysis beyond that using Avida’s Analyze Mode was performed using the Python packages NumPy version 1.12.1 [43], SciPy version 0.19.0 [44], and Pandas version 0.20.1 [45]; figures were generated using the Python package Matplotlib version 2.0.2 [46].

## Results

### Large genome size increases the extinction risk of small populations

To test if genome expansions and large genome size enhanced the probability of population extinction, we evolved populations across a range of sizes at both high (1.5 mutations/genome/generation) and low (0.15 mutations/genome/generation) with both a fixed genome size and an evolving genome size. Under the low mutation rate regime, populations with variable genome sizes had greater rates of extinction than those with fixed small genomes (Fig. 1a). Under the high mutation rate regime, there was no significant difference between populations with a variable genome size and populations with a constant genome size (Fig. 1a). Estimations of the time to extinction further support these trends. In the low mutation regime, populations where genome size could evolve went extinct in fewer generations than those where genome size was constant; there were no differences in the high mutation rate regime (Fig. 1b).

**Figure 1:**
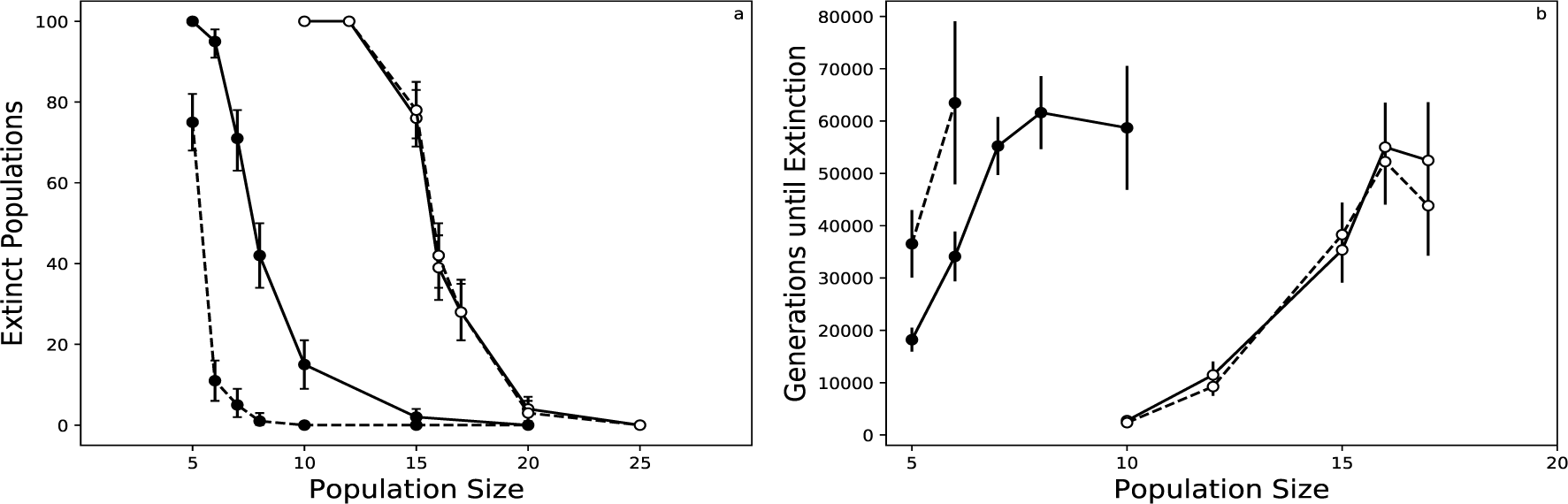
Possibility of genome expansions increase extinction in low mutation rate populations. a) Number of extinct populations as a function of population size. Full lines represent variable genome size populations; dashed lines represent fixed genome size populations. Black circles represent low mutation rate populations; white circles represent high mutation rate populations. Error bars are bootstrapped 95% confidence intervals (10^4^ samples). b) Time to extinction for population size and mutation rate combinations where at least ten populations went extinct. Lines and colors same as in panel a. Error bars represent 2× standard error of the mean. Data only shown for those treatments that resulted in at least ten extinct populations.

Next, we compared the final evolved genome size between genotypes from extinct populations and surviving populations. Across the range of population sizes for which at least 10 populations both survived and went extinct, “extinct” genotypes evolved larger genomes than those “surviving” genotypes in the low mutation rate regime (Fig. 2a). In the high mutation rate regime, one population size (*N* = 15 individuals) led to surviving populations evolving larger genomes, while there was no statistically-significant difference for the other population sizes (Fig. 2b). Together, these results suggest that genome expansions and large genome size can enhance the risk of small population extinction if the initial mutation rate is too low for extinction to occur. Now, we will focus on examining the mechanism behind the relationship between genome size and extinction in the low mutation rate populations.

**Figure 2:**
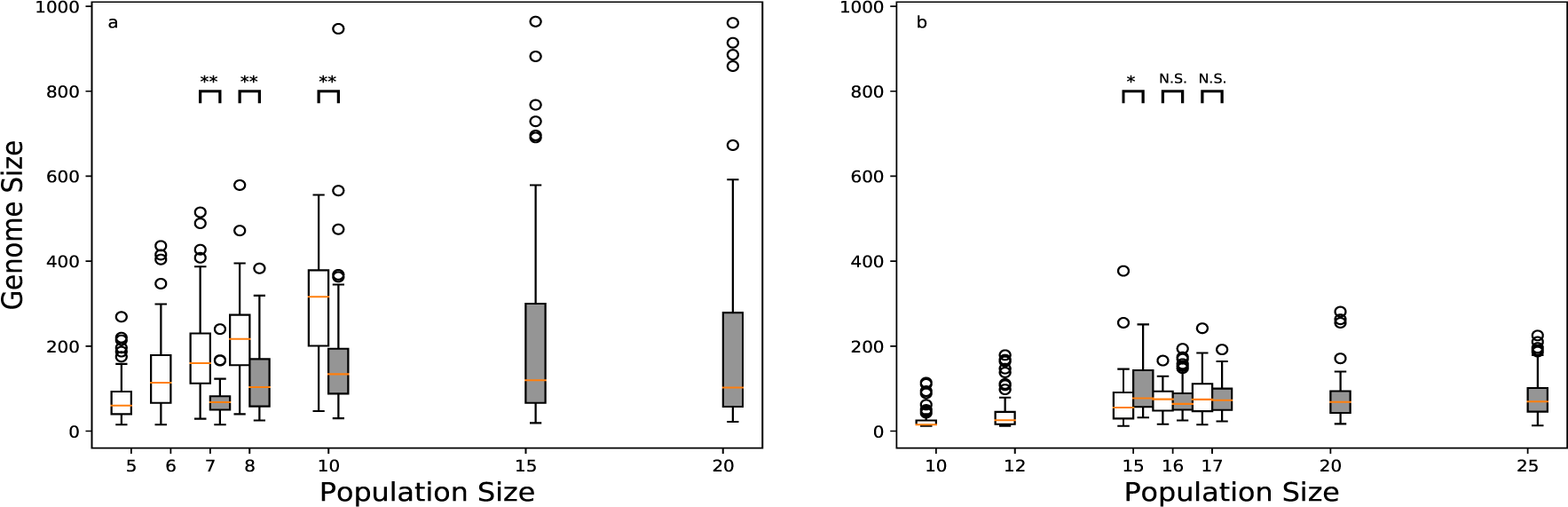
Extinct populations evolved larger genomes. a) Final genome size for the low mutation rate populations as a function of population size. Populations that survived are shown with gray boxplots; populations that went extinct are shown with white boxplots. Red bars are the median value, boxes are the first and third quartile, upper/lower whiskers extend up to the 1.5 times the interquartile range, and circles are outliers. ** indicates MannWhitney U *p* < 10^−4^, * indicates *p* < 10^−2^, and N.S. indicates *p* > 0.05. Population sizes where fewer than ten populations went extinct (or survived) not shown. b) Final genome size for the high mutation rate populations as a function of population size. Description same as in panel a. Population sizes where fewer than ten populations went extinct (or survived) not shown.

#### Extinction and large genome size is associated with increases in the lethal mutational load

In a constant environment with soft selection, Avidian populations only face population-size reductions through one mechanism: parent avidians produce non-viable (or infertile) offspring that replace viable avidians. In other words, the lethal mutational load should drive population reduction and eventually population extinction. It is therefore possible that the increased genomic mutation rate that co-occurs with genome size increases specifically increased the genomic lethal mutation rate and that led to increased rates of population extinction. We first tested whether larger genomes had an increased lethal mutation rate. Genome size was correlated with the lethal mutation rate across genotypes from all population sizes, supporting the hypothesis that increases in genome size result in increased lethal mutational load and eventually population (Fig. 3a; Spearman’s *ρ* ≈ 0.75,*p* = 1.77 × 10^−148^). Next, we examined whether populations that went extinct had previously evolved greater lethal mutation rates than surviving populations. As with the trend for genome size, extinct populations evolved greater lethal mutations rates than surviving populations (Fig. 3b).

**Figure 3:**
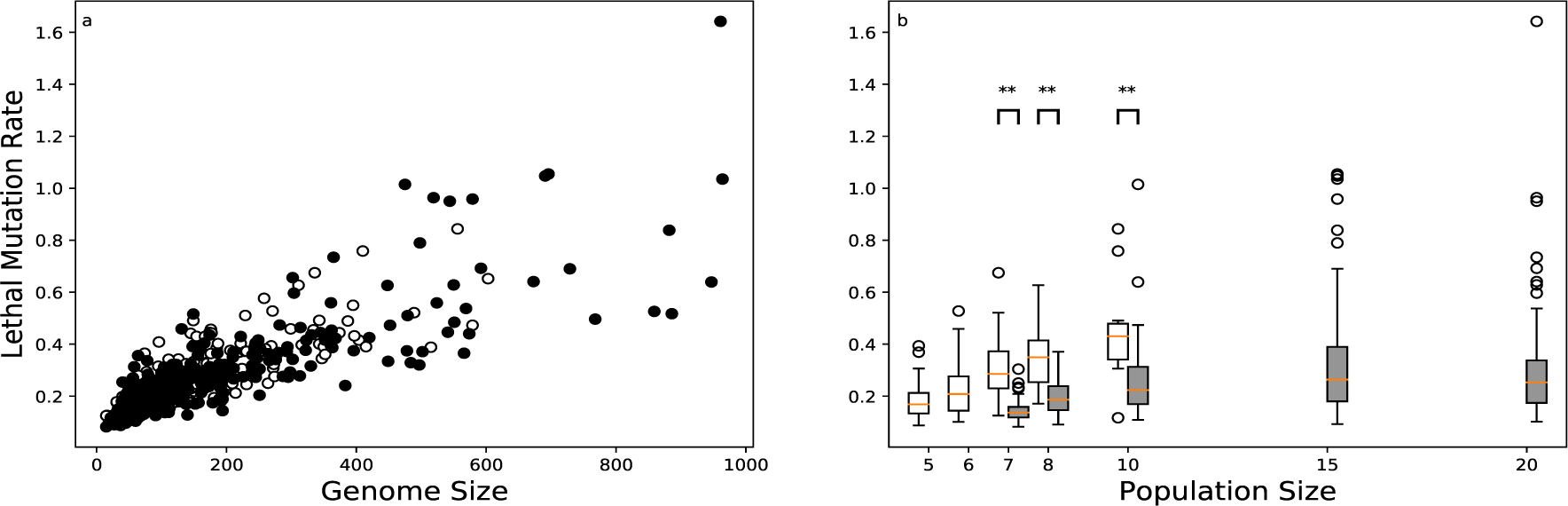
Lethal mutation rate correlates with genome size and population extinction. a) The lethal mutation rate as a function of genome size for the final genotypes from each evolved low mutation rate population. White circles are extinct populations; gray circles are surviving populations. b) The lethal mutation rate for extinct and surviving populations across population sizes. Boxplots as previously described. Colors and significance symbols same as Figure 2. Population sizes where fewer than ten populations went extinct (or survived) not shown.

The previous data support the hypothesis that genome expansions drive population extinction by increases in the lethal mutation rate and thus the lethal mutational load. However, it is unclear whether genome expansions themselves increase the likelihood of lethal mutations or whether genome expansions merely potentiate the ability for further evolution to increases the lethal mutation rate. To test these two scenarios, we examined the evolutionary histories (i.e., line-of-descents) for all 100 *N* = 20 low mutation-rate populations. We then examined the relationship between changes in genome size and changes in the lethal mutation rate (Fig. 4a). When genome size was constant, the lethal mutation rate did not change on average (mean change = 7 × 10^−4^, 95% Confidence Interval = ±2 × 10^−3^). Genome size increases on average increased the lethal mutation rate (mean change = 0.024, 95% Confidence Interval = ±0.0023), while genome size decreases on average decreased the lethal mutation rate (mean change = −0.033, 95% Confidence Interval = ±0.0040). Additionally, the change in genome size positively correlates with the change in the lethal mutation rate (Fig. 4b; Spearman’s *ρ* = 0.67, *p* ≈ 0.0).

**Figure 4:**
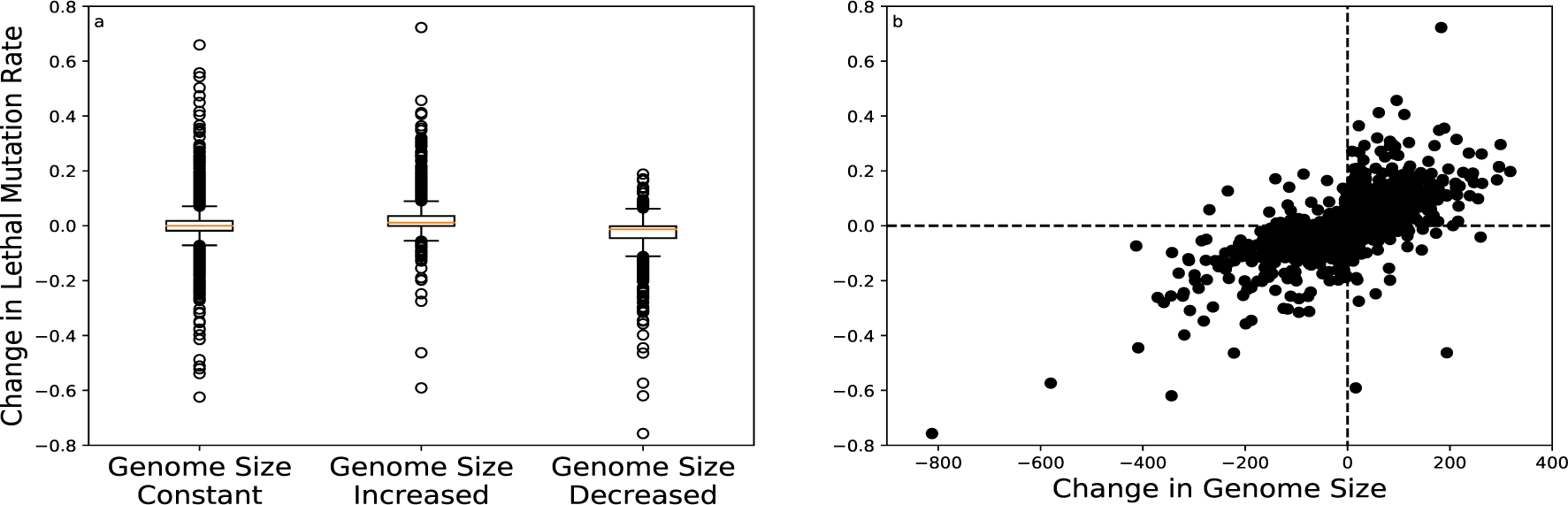
Insertions and deletions directly change the lethal mutation rate. a) Change in the lethal mutation rate as a function of a mutation’s effect on genome size. Boxplots as previously described. Each data point represents the a genotype from a reduced evolutionary lineage of a population with 20 individuals. b) Relationship between a mutation’s change in genome size and the change in the lethal mutation rate. Data same as in panel a. Dashed line represent no change. Data points comparing genotypes with equal genome size were excluded.

#### Lethal mutation rate and stochastic viability drive population extinction

Finally, to establish the role of the lethal mutation rate in driving population extinction, we performed additional evolution experiments to test whether the prevention of lethal mutations would prevent population extinction. We repeated our initial experiments (Fig. 1), except offspring with lethal mutations were reverted to their parental genome (lethal-reversion treatment; see Methods for details). We also did the same experiment where deleterious, but non-lethal, mutations were reverted in order to test if deleterious mutations contributed to extinction. When populations evolved without deleterious mutations, extinction rates were similar to, if not greater than, those for populations that evolved with deleterious mutations (Fig. 5a). Populations that evolved with fixed-size genomes and without lethal mutations never went extinct, demonstrating how the lack of lethal mutations can prevent extinction (Fig. 5b). However, when these populations evolved with variable genome sizes, extinction still occurred, although at a lower rate than when lethal mutations were present (Fig. 5b).

**Figure 5:**
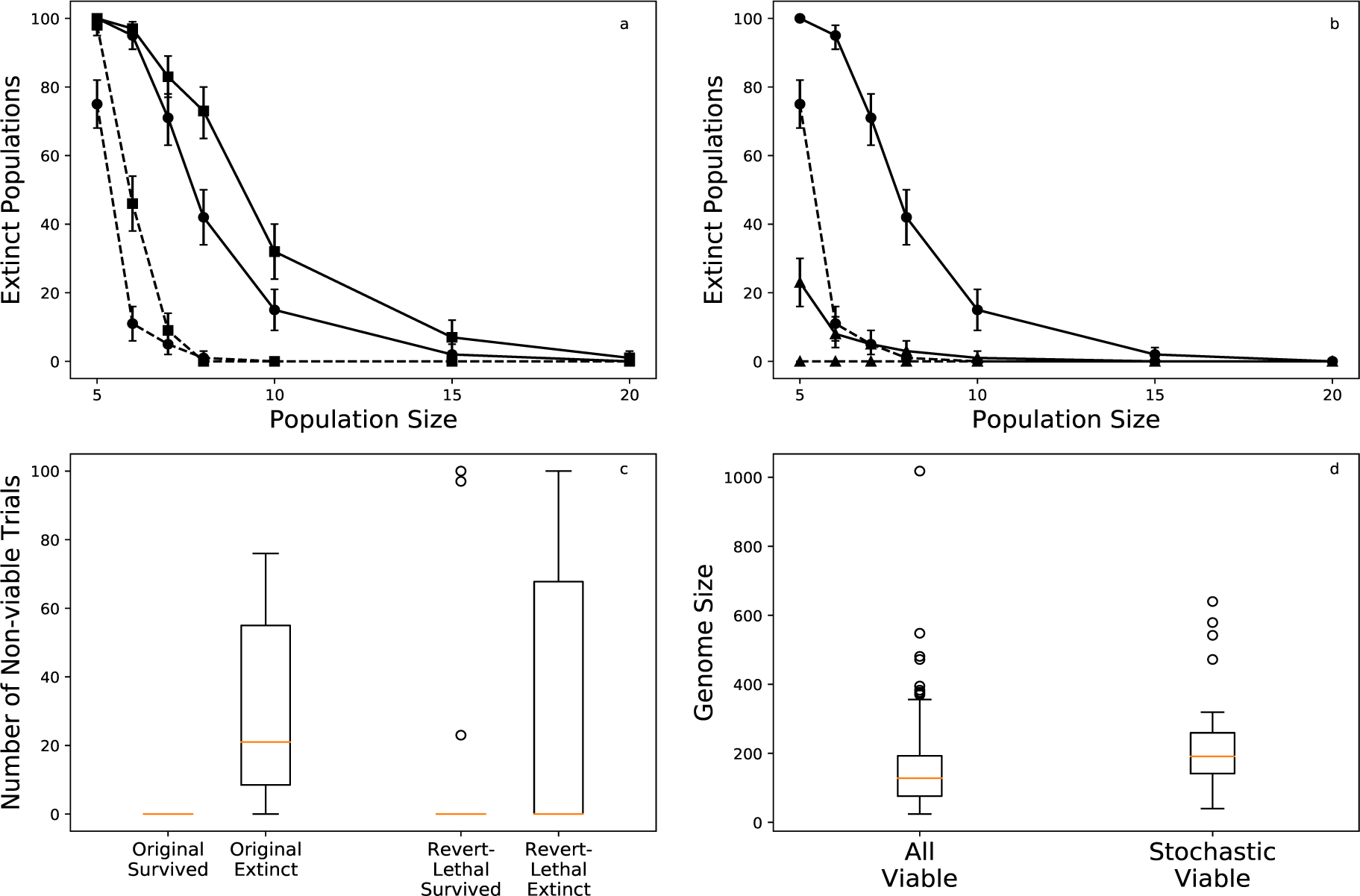
Evolution of stochastic viability contributes to extinction risk. a) Number of population extinctions (out of 100 replicates) as a function of population size. Squares represent deleterious-reversion populations and circles represent populations where lethal mutations were not reverted. Dashed lines represent populations with fixed-size genomes; solid lines represent populations with variable-size genomes. Error bars are 95% confidence intervals generated using bootstrap sampling (10^4^ samples). b) Number of population extinctions (out of 100 replicates) as a function of population size. Triangles represent lethal-reversion populations. All other symbols same as in panel a. c) Number of viability trials (out of 100) that estimated that a given genotype was not viable. Values between 0 and 100 indicate stochastic viability. “Original” refers to the 100 genotypes from the populations of 8 individuals that evolved with lethal mutations. “Revert-lethal” refers the 100 genotypes from the *N* = 5 lethal-reversion populations. Boxplots as previously described. d) Genome size as a function of whether a genotype was measured as stochastic-viable or deterministicviable. Data are the 100 genotypes from the *N* = 5 lethal-reversion populations. Boxplots as previously described.

While these data demonstrate that lethal mutations do primarily drive extinction risk, they also show that there is a second factor that relates genome size to extinction. This is surprising, as lethal mutations are the only direct mechanism to cause extinction in Avida. One possible explanation for extinction in the lethal-reversion populations is that mutants arise in these populations that are initially viable, but later become non-viable. In other words, these populations evolve stochastic viability, where characteristics of the random numbers the avidians input during their life-cycle determine their ability to reproduce. These genotypes with stochastic viability would, on occasion, not be measured as lethal mutants, and thus enter the population even when lethal mutations are reverted. As they reproduce, these stochastic-viable genotypes will input other numbers and thus become, in effect, a lethal mutation and subsequently lead to population extinction. To test if these populations that went extinct without lethal mutations did evolve stochastic viability, we tested the viability of all 100 genotypes from the variable genome size *N* = 5 populations; each genotype was tested 100 times. We also performed the same tests with the 100 genotypes from the *N* = 8 populations that evolved with lethal mutations to see if these mutants arose in our original populations.

For both sets of genotypes, we found that some genotypes were stochastically viable (Fig. 5c). Of the 23 genotypes from populations that went extinct in the lethal-reversion treatment, 19 displayed stochastic viability. The remaining four populations did not show signs of stochastic viability; however, as we only sampled one genotype per population ten generations before extinction, there was likely stochastic-viable replicators among the remaining individuals in the populations. No genotypes from surviving populations were stochastically-viable. Of the 43 genotypes from populations that went extinct among our original treatment genotypes, eight displayed stochastic viability. Two genotypes from surviving populations were stochastically-viable, suggesting these populations would have an enhanced likelihood of extinction if the experiment had continued. Finally, we compared the genome sizes between genotypes from the lethal-reversion genotypes that were always measured as viable and those that measured as stochastic-viable. Stochastic-viable genotypes evolved larger genomes than deterministic-viable genotypes (Mann-Whitney U, median = 128 instructions vs. median = 191 instructions, *p* < 2 × 10^−4^; Fig. 5d), further suggesting that increased genome size can lead to the evolution of stochastic viability and eventually population extinction.

### Discussion

We explored the role of genome size in the extinction of small populations and found that low mutation rate populations with variable genome sizes go extinct at a higher rate than those with a fixed genome size. Large genome size enhances the rate of small population extinction due to two factors: increased lethal mutation rates and increased appearance of genotypes with stochastic viability. Extinction occurs because, as genome size and the lethal mutation rate increases, the population-level lethal mutational load increases, driving population collapse. These increases in the lethal mutation rate are directly driven by genome expansion mutations, not by evolution after genome expansion, while deletions lowered the lethal mutation rate. Finally, we showed populations with large genomes are at risk of evolving genotypes with stochastic viability and this increases the contribution of large genome size to extinction.

The most prominent model of small population extinction is the mutational meltdown model [15, 17, 19], which argues that even intermediate-sized asexual and sexual populations (i.e., 10^3^ individuals) can go extinct on the order of thousands of generations. It is worth comparing our results from the predictions of the meltdown model, namely that only very small populations go extinct in Avida, and extinction occurs on a much longer timescale than in the mutational meltdown model. The contrast between our results and previous results are likely due to differences in the character of selection between the two models. Selection is hard in mutational meltdown models, and the accumulation of deleterious mutations due to genetic drift reduces the probability of offspring surviving to reproduce [15]. In other words, the accumulation of deleterious mutations directly increases the probability that offspring will be non-viable. In Avida, selection on deleterious mutations is soft; accumulation of deleterious mutations due to drift is unrelated to viability and the likelihood of offspring receiving lethal mutations. The lethal mutation rate will only increase indirectly in Avida due to accumulation of genome expansions in very small populations. Without the positive feedback loop between deleterious mutation accumulation and population size, avidian populations only evolve a high rate of non-viabile mutants if they undergo large genome expansions, thus explaining the trends we saw here.

These differences between extinction in hard selection models and the hybrid selection model we used here emphasizes the need to consider whether selection in biological populations is primarily hard or soft. Unfortunately, there has been little resolution on this question [47, 48]. There is some evidence that soft selection may be more relevant to evolutionary dynamics than hard selection. For instance, soft selection has been invoked as an explanation for why humans are able to experience high rates of deleterious mutations per generation [49, 50]. Moreover, the persistence of small, isolated populations [51, 52, 53] suggests that not only is selection primarily soft in nature, but that the extinction dynamics we study here are relevant to a subset of biological populations. While large genome size may not be the factor that causes populations to begin their ecological decline, it could drive an already-reduced population to extinction.

We discovered two genetic mechanisms responsible for the relationship between large genome size and small population extinction. First, genome expansions directly increase the likelihood that an offspring will receive a lethal mutation and thus be non-viable. This is contrary to expectations from biological organisms, where it is often assumed that genome expansions, i.e., gene/genome duplications, increase robustness and reduce the lethality of mutations by multiplying the copies of essential genes [54]. However, recent research has shown that gene duplication can also result in increased mutational fragility, not just mutational robustness [55]. Additionally, one major component of genome growth in many multicellular eukaryotes is the expansion of transposable elements [56]. Any increase in the number of active transposable elements in a genome should increase the lethal mutation rate, as we have shown here, by increasing the likelihood that any essential gene will be mutated. Selfish genetic elements such as transposons were the original proposed mechanism to explain the relationship between genome size and extinction risk in plant species [23]. However, there is no equivalent to transposable elements in Avida, so we cannot directly test relevance of this mechanism here.

Our second proposed mechanism underlying the connection between genome size and population extinction is the evolution of stochastic viability, or genotypes that could only reproduce under some environmental conditions (i.e., random number inputs). The connection behind stochastic viability and extinction in small populations is intuitive. Mutations causing stochastic viability likely have a weak effect (due to their stochastic nature) and can fix in small populations due to weakened selection. After fixation, the lethality of these mutation may be stochastically revealed, and extinction occurs. However, studies on the functional consequences of mutations responsible for extinction are novel [57, 58] and it is uncertain whether these mutations arise in populations at high extinction risk. One suggestion that mutations with stochastic effects might be relevant to population extinction comes from microbial experimental evolution. It has been shown that small populations have reduced extinction risk if they over-express genes encoding molecular chaperones that assist with protein folding [59]. These over-expressed chaperones presumably compensate for other mutations that cause increased rates of stochastic protein misfolding. Therefore, mutations responsible for an increased likelihood of protein misfolding may be an example of a class of mutations with a stochastic effect that enhance extinction risk. However, this is only speculation and further work is needed to see if stochastic viability is a possible mechanism behind extinction risk.

In a previous study, we observed that small populations evolved the largest genomes, the greatest phenotypic complexity, and the greatest rates of extinction [31]. This result raised the question of whether greater biological complexity itself could increase a population’s rate of extinction, a question we explored here. Although we did not test whether increased phenotypic complexity had a role in extinction, we have shown that genomic complexity, measured in terms of genome size, did drive small-population extinction. While it is possible that phenotypic complexity also enhanced the likelihood of extinction, the Avida phenotypic traits likely do not increase the lethal mutation rate. Thus, both high extinction rates and increased phenotypic complexity arise due to the same mechanism: greater genome size. This result illustrates an evolutionary constraint for small populations. While weakened selection and stronger genetic drift can lead to increases in biological complexity, small populations must also evolve genetic architecture that reduces the risk of extinction. Otherwise, small populations cannot maintain greater complexity and their lethal mutational load drives them to extinction.

## Data Accessibility

The Avida software is available for free at (https://github.com/devosoft/avida). All Avida scripts, data analysis scripts, and data used to generate the figures are available at Dryad

## Author Contributions

T.L. and C.A. designed the study, wrote the manuscript, and approved the final version. T.L. performed the experiments and data analysis.

## Acknowledgments

We thank M. Zettlemoyer and C. Weisman for comments on drafts of the manuscript. T.L. acknowledges a Michigan State University Distinguished Fellowship, a BEACON fellowship, and the Russell B. Duvall award for support. This work was supported in part by Michigan State University through computational resources provided by the Institute for Cyber-Enabled Research.

## References

[1] Jablonski D. Background and mass extinctions: the alternation of macroevolutionary regimes. Science. 1986;231(4734):129–133.

[2] Barnosky AD, Matzke N, Tomiya S, Wogan GO, Swartz B, Quental TB, et al. Has the Earth’s sixth mass extinction already arrived? Nature. 2011;471(7336):51–57.

[3] Maynard Smith J. The causes of extinction. Philosophical Transactions of the Royal Society of London B: Biological Sciences. 1989;325(1228):241–252.

[4] Spielman D, Brook BW, Frankham R. Most species are not driven to extinction before genetic factors impact them. Proceedings of the National Academy of Sciences. 2004;101(42):15261–15264.

[5] O’Grady JJ, Brook BW, Reed DH, Ballou JD, Tonkyn DW, Frankham R. Realistic levels of inbreeding depression strongly affect extinction risk in wild populations. Biological Conservation. 2006;133(1):42–51.

[6] Matthies D, Bräuer I, Maibom W, Tscharntke T. Population size and the risk of local extinction: empirical evidence from rare plants. Oikos. 2004;105(3):481–488.

[7] Melbourne BA, Hastings A. Extinction risk depends strongly on factors contributing to stochasticity. Nature. 2008;454(7200):100–103.

[8] Lindsey HA, Gallie J, Taylor S, Kerr B. Evolutionary rescue from extinction is contingent on a lower rate of environmental change. Nature. 2013;494(7438):463–467.

[9] Urban MC. Accelerating extinction risk from climate change. Science. 2015;348(6234):571–573.

[10] Clavero M, García-Berthou E. Invasive species are a leading cause of animal extinctions. Trends in Ecology and Evolution. 2005;20(3):110–110.

[11] Pedersen AB, Jones KE, Nunn CL, Altizer S. Infectious diseases and extinction risk in wild mammals. Conservation Biology. 2007;21(5):1269–1279.

[12] Dunn RR, Harris NC, Colwell RK, Koh LP, Sodhi NS. The sixth mass coextinction: are most endangered species parasites and mutualists? Proceedings of the Royal Society of London B: Biological Sciences. 2009;276(1670):3037–3045.

[13] Raup DM. Extinction: bad genes or bad luck? New York: WW Norton & Company; 1992.

[14] Turner CB, Blount ZD, Lenski RE. Replaying Evolution to Test the Cause of Extinction of One Ecotype in an Experimentally Evolved Population. PloS One. 2015;10(11):e0142050.

[15] Lynch M, Gabriel W. Mutation load and the survival of small populations. Evolution. 1990;44(7):1725–1737.

[16] Whitlock MC, Bürger R, Dieckmann U. Fixation of new mutations in small populations. In: Ferriere R, Dieckmann U, Couvet D, editors. Evolutionary Conservation Biology. Cambridge: Cambridge University Press; 2004. p. 155–170.

[17] Lynch M, Bürger R, Butcher D, Gabriel W. The mutational meltdown in asexual populations. Journal of Heredity. 1993;84(5):339–344.

[18] Gabriel W, Lynch M, Burger R. Muller’s ratchet and mutational meltdowns. Evolution. 1993;47(6):1744–1757.

[19] Lynch M, Conery J, Burger R. Mutation accumulation and the extinction of small populations. American Naturalist. 1995;146(4):489–518.

[20] Lande R. Risk of population extinction from fixation of new deleterious mutations. Evolution. 1994;48(5):1460–1469.

[21] Zeyl C, Mizesko M, De Visser J. Mutational meltdown in laboratory yeast populations. Evolution. 2001;55(5):909–917.

[22] Singh T, Hyun M, Sniegowski P. Evolution of mutation rates in hypermutable populations of Escherichia coli propagated at very small effective population size. Biology Letters. 2017;13(3):20160849.

[23] Vinogradov AE. Selfish DNA is maladaptive: evidence from the plant Red List. Trends in Genetics. 2003;19(11):609–614.

[24] Vinogradov AE. Genome size and extinction risk in vertebrates. Proceedings of the Royal Society of London B: Biological Sciences. 2004;271:1701–1706.

[25] Hindré T, Knibbe C, Beslon G, Schneider D. New insights into bacterial adaptation through *in vivo* and *in silico* experimental evolution. Nature Reviews Microbiology. 2012;10(5):352–365.

[26] Kawecki TJ, Lenski RE, Ebert D, Hollis B, Olivieri I, Whitlock MC. Experimental evolution. Trends in Ecology & Evolution. 2012;27(10):547–560.

[27] Batut B, Parsons DP, Fischer S, Beslon G, Knibbe C. *In silico* experimental evolution: a tool to test evolutionary scenarios. BMC Bioinformatics. 2013;14(Suppl 15):S11.

[28] Adami C. Digital genetics: Unravelling the genetic basis of evolution. Nature Reviews Genetics. 2006;7(2):109–118.

[29] Pennock RT. Models, simulations, instantiations, and evidence: the case of digital evolution. Journal of Experimental & Theoretical Artificial Intelligence. 2007;19(1):29–42.

[30] Ofria C, Bryson DM, Wilke CO. Avida: A Software Platform for Research in Computational Evolutionary Biology. In: Maciej Komosinski AA, editor. Artificial Life Models in Software. Springer London; 2009. p. 3–35.

[31] LaBar T, Adami C. Different evolutionary paths to complexity for small and large populations of digital organisms. PLoS Computational Biology. 2016;12(12):e1005066.

[32] Adami C. Introduction to Artificial Life. TELOS, Springer Verlag; 1998.

[33] Gupta A, LaBar T, Miyagi M, Adami C. Evolution of Genome Size in Asexual Digital Organisms. Scientific Reports. 2016;6:25786.

[34] Misevic D, Lenski RE, Ofria C. Sexual reproduction and Muller’s ratchet in digital organisms. In: Ninth International Conference on Artificial Life; 2004. p. 340–345.

[35] Elena SF, Wilke CO, Ofria C, Lenski RE. Effects of population size and mutation rate on the evolution of mutational robustness. Evolution. 2007;61(3):666–674.

[36] LaBar T, Adami C. Evolution of Drift Robustness in Small Populations. Nature Communications. 2017;8:1012.

[37] Yedid G, Ofria C, Lenski R. Historical and contingent factors affect re-evolution of a complex feature lost during mass extinction in communities of digital organisms. Journal of Evolutionary Biology. 2008;21(5):1335–1357.

[38] Yedid G, Ofria CA, Lenski RE. Selective press extinctions, but not random pulse extinctions, cause delayed ecological recovery in communities of digital organisms. The American Naturalist. 2009;173(4):E139–E154.

[39] Yedid G, Stredwick J, Ofria CA, Agapow PM. A comparison of the effects of random and selective mass extinctions on erosion of evolutionary history in communities of digital organisms. PloS One. 2012;7(5):e37233.

[40] Strona G, Lafferty KD. Environmental change makes robust ecological networks fragile. Nature Communications. 2016;7:12462.

[41] Lenski RE, Ofria C, Pennock RT, Adami C. The evolutionary origin of complex features. Nature. 2003;423(6936):139–144.

[42] Covert AW, Lenski RE, Wilke CO, Ofria C. Experiments on the role of deleterious mutations as stepping stones in adaptive evolution. Proceedings of the National Academy of Sciences. 2013;110(34):E3171–E3178.

[43] Walt Svd, Colbert SC, Varoquaux G. The NumPy array: a structure for efficient numerical computation. Computing in Science & Engineering. 2011;13(2):22–30.

[44] Jones E, Oliphant T, Peterson P, et al.. SciPy: Open source scientific tools for Python; 2001—. Available from: http://www.scipy.org/.

[45] McKinney W, et al. Data structures for statistical computing in Python. In: Proceedings of the 9th Python in Science Conference. vol. 445. SciPy Austin, TX; 2010. p. 51–56.

[46] Hunter JD. Matplotlib: A 2D graphics environment. Computing In Science & Engineering. 2007;9(3):90–95.

[47] Wallace B. Fifty Years of Genetic Load: An Odyssey. Cornell University Press; 1991.

[48] Reznick D. Hard and soft selection revisited: How evolution by natural selection works in the real world. Journal of Heredity. 2015;107(1):3–14.

[49] Lesecque Y, Keightley PD, Eyre-Walker A. A resolution of the mutation load paradox in humans. Genetics. 2012;191(4):1321–1330.

[50] Charlesworth B. Why we are not dead one hundred times over. Evolution. 2013;67(11):3354–3361.

[51] Xue Y, Prado-Martinez J, Sudmant PH, Narasimhan V, Ayub Q, Szpak M, et al. Mountain gorilla genomes reveal the impact of long-term population decline and inbreeding. Science. 2015;348(6231):242–245.

[52] Robinson JA, Ortega-Del Vecchyo D, Fan Z, Kim BY, Marsden CD, Lohmueller KE, et al. Genomic flatlining in the endangered island fox. Current Biology. 2016;26(9):1183–1189.

[53] Benazzo A, Trucchi E, Cahill JA, Delser PM, Mona S, Fumagalli M, et al. Survival and divergence in a small group: The extraordinary genomic history of the endangered Apennine brown bear stragglers. Proceedings of the National Academy of Sciences. 2017;114(45):E9589–E9597.

[54] Gu Z, Steinmetz LM, Gu X, Scharfe C, Davis RW, Li WH. Role of duplicate genes in genetic robustness against null mutations. Nature. 2003;421(6918):63–66.

[55] Diss G, Gagnon-Arsenault I, Dion-Coté AM, Vignaud H, Ascencio DI, Berger CM, et al. Gene duplication can impart fragility, not robustness, in the yeast protein interaction network. Science. 2017;355(6325):630–634.

[56] Kidwell MG. Transposable elements and the evolution of genome size in eukaryotes. Genetica. 2002;115(1):49–63.

[57] Rogers RL, Slatkin M. Excess of genomic defects in a woolly mammoth on Wrangel island. PLoS Genetics. 2017;13(3):e1006601.

[58] Fry E, Kim SK, Chigurapti S, Mika KM, Ratan A, Dammermann A, et al. Accumulation And Functional Architecture Of Deleterious Genetic Variants During The Extinction Of Wrangel Island Mammoths. bioRxiv. 2017;p. 137455.

[59] Sabater-Muñoz B, Prats-Escriche M, Montagud-Martínez R, López-Cerdán A, Toft C, Aguilar-Rodríguez J, et al. Fitness trade-offs determine the role of the molecular chaperonin GroEL in buffering mutations. Molecular biology and evolution. 2015;32(10):2681–2693.

